# bL9 prevents recoding in collided ribosomes

**DOI:** 10.64898/2026.11.03.763934

**Authors:** Rebecca E. Zordan, Jasmine L. James, Rachel Green, Allen R. Buskirk

## Abstract

When a ribosome stalls and an upstream ribosome collides with it, the resulting interface recruits factors that initiate ribosome rescue pathways. Recent cryo-EM structures of collided ribosomes from bacteria reveal that bL9 from the leading ribosome reaches back to contact the 30S subunit of the trailing ribosome. The role of bL9 in translation is poorly understood; while bL9 mutants show high levels of recoding (i.e. frameshifting and hopping), the mechanism has remained unclear. Here we show that bL9 functions in the context of ribosome collisions: loss of bL9 destabilizes the interaction between collided ribosomes and sensitizes cells to antibiotics that induce collisions such as erythromycin and fusidic acid. Moreover, luciferase reporters reveal that frameshifting and hopping levels increase in strains lacking bL9 (or with point mutants in bL9 at the collision interface) but only in the context of ribosome collisions. These findings demonstrate that in wild-type bacterial ribosomes, bL9 prevents frameshifting, hopping, and other deleterious recoding events following collisions.

## Introduction

During protein synthesis, translating ribosomes stall due to mRNA damage (1), stretches of rare codons that are slowly decoded (2, 3), or peptide sequences that inhibit their own translation (4). In bacteria, antibiotics also pose a significant threat, stalling elongating ribosomes through many mechanisms (5, 6). Ribosome rescue factors mitigate the negative consequences of ribosome stalling by recycling the ribosomal subunits, targeting the nascent peptide for degradation, and degrading the problematic mRNA, as reviewed in (7).

Over the last decade, a consensus has emerged first from work in yeast and mammalian cells (8, 9), and more recently in bacteria (10, 11), that ribosome collisions are the critical trigger of ribosome rescue pathways, allowing rescue factors to discriminate between stalled ribosomes and actively translating ones. In *E. coli*, for example, collisions recruit the endonuclease SmrB, which binds to ribosomal proteins on both sides of the collision interface, positioning its active site to promote cleavage and rapid decay of the problematic transcript (10). Another rescue factor, the ATPase HrpA, binds to mRNA downstream of the leading ribosome with its helicase domain and senses the collided ribosome using its C-terminal domain (12). HrpA then uses ATP hydrolysis activity to destabilize the leading ribosome, splitting it into subunits and allowing the trailing ribosome to continue translation (12). The activity of both of these rescue factors is dependent on ribosome collisions, where the interface between the ribosomes serves as a signal that something has gone wrong in translation (7).

Cryo-EM structures of collided disomes from bacteria reveal that the collision interface is formed almost entirely by the juxtaposition of the small subunits of the two ribosomes (Figure 1A) (10, 11). In many ways the overall architecture of the collided disomes in bacteria resembles structures from yeast and mammalian disomes, even if the details of the interactions between the ribosomes differ slightly (9, 10, 13). Yet there is one major difference: in bacterial disomes, the bL9 protein (unique to bacteria) reaches back from the large subunit of the leading ribosome to make significant contacts with 16S rRNA in the trailing ribosome (Figure 1A). This bridging interaction of bL9 has been reported in cryo-EM studies of disomes from *E. coli* and *B. subtilis* (10, 11, 14, 15), cryo-electron tomography studies of *M. pneumoniae* (16, 17), and even in x-ray crystallography structures of 70S monosomes from thermophilic bacteria, where the bL9 interaction between monosomes formed part of the crystal lattice (18–20).

**Figure 1.**
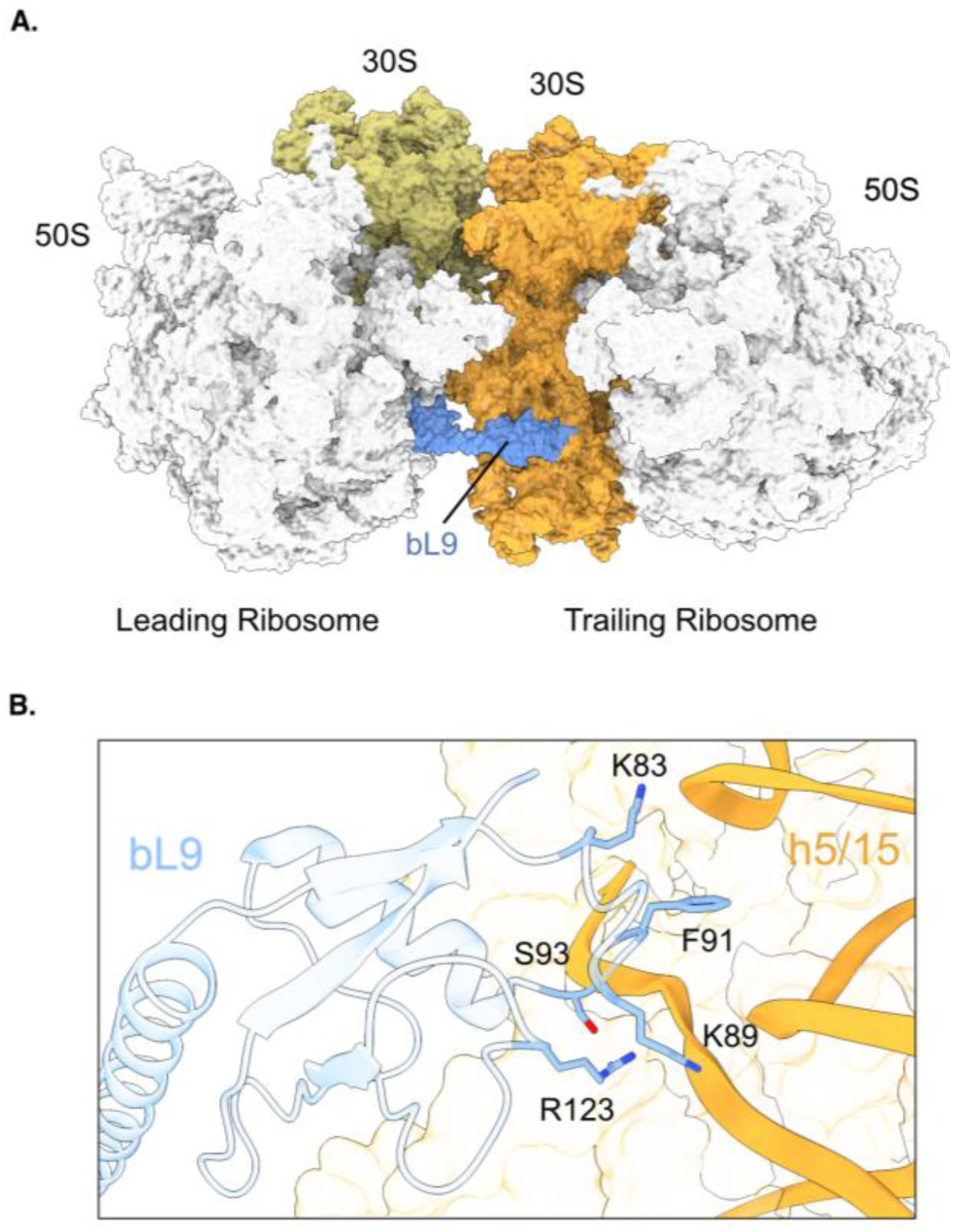
bL9 bridges collided ribosomes. **(A)** A cryo-EM structure of collided *E. coli* ribosomes (PDBID: 8PKL & 8PEG) shows the bL9 of the leading ribosome (blue) making contacts with the small subunit of the trailing ribosome (dark orange). **(B)** The inset shows the side chains of key bL9 residues from the leading ribosome, and how they interact with the h5/h15 helix of 16S rRNA in the small subunit of the trailing ribosome.

bL9 has two domains linked by an α-helix; the N-terminal domain binds the large subunit on the surface near the E site. While the C-terminal domain binds near L2 and bS6 in cryo-EM structures of 70S monosomes (21), in the context of collisions it reaches out to the trailing ribosome, where it binds the 16S rRNA near helices h5 and h15 (10, 15). Studies of *ex vivo* polysomes and *in vitro* translation reactions led to a high-resolution view of this interface, where the conserved residue Phe91 forms a stacking interaction with both G57 and U368 of the 16S rRNA in the trailing ribosome, and where three positively charged residues interact with the sugar-phosphate backbone (Figure 1B) (15). While the significance of the bL9 bridge is unknown, it is tempting to speculate that it would block binding of elongation factors EFG and EFTu to the trailing ribosome, preventing futile attempts at elongation. Cryo-ET structures of translation in *M. pneumoniae* reveal several distinct disome structures with the ribosomes separated by 4.0, 2.8, and 2.2 nm; the disomes separated by 2.2 nm closely resemble the *in vitro* collided disomes structures bound to ribosome rescue factors SmrB and HrpA (17). This observation raises the possibility that bL9 may obstruct the trailing ribosome before the tight collision interface forms and ribosome rescue factors abort protein synthesis.

Two phenotypes of strains with mutant bL9 have been puzzling for many years. First, rather unusually for a ribosomal protein, bL9 is not essential for translation; knockouts are viable and show only a minor growth defect (22). Several genome-wide screens found that strains lacking bL9 are hypersensitive to antibiotics that target elongating ribosomes, particularly fusidic acid (23–25). This sensitivity is difficult to explain in the context of a single 70S monosome given that fusidic acid traps EFG on the ribosome far away from bL9 (26, 27). Secondly, a mutation in bL9 (Ser93 to Phe) was isolated in a genetic selection for increased levels of bypassing in *gene 60* in T4 phage (28). In this remarkable recoding event, when ribosomes reach a premature stop codon in *gene 60*, the peptidyl-tRNA dissociates from the mRNA, and the ribosome slides 50 nt down the mRNA until the peptidyl-tRNA pairs with a cognate codon and translation resumes, completing translation of *gene 60* (29, 30). While this mechanism has been studied for nearly forty years (31–34), it has remained mysterious how bL9 somehow plays a role in suppressing bypassing. The fact that the S93F mutation lies at the collision interface (Figure 1B) raises the possibility that the role of bL9 in suppressing recoding is in the context of ribosome collisions.

The increase in recoding in strains with mutated bL9 is not specific to *gene 60*: high levels of recoding have also been reported in the endogenous *ybeL* gene in strains lacking bL9 (35). This recoding happens when a ribosome stalls at the stop codon in the *ybeL* gene, whose final residues, Glu-Pro, inhibit termination (36). These two C-terminal residues are sufficient for stalling, presumably due to the steric constraints of the Pro residue. Using pre-steady state kinetics in a reconstituted system, we previously reported that peptidyl hydrolysis following Met-Glu-Pro is 275-fold slower than Met-Glu-Phe (37). Hayes *et al.* carefully characterized the mechanisms by which ribosome stalling at the stop codon in *ybeL* leads to +1 frameshifting and hopping, depending on the codon context (35).

We note that there is a previous report claiming that ribosome collisions affect frameshifting efficiency on reporters containing programmed frameshifting sites (38). It is unlikely that ribosomes stall at these slippery sequences. The lack of evidence for collisions and the inconsistent findings raise doubts about the conclusions of this study.

Here, we report experiments designed to test the role of the bL9 bridge between collided disomes. We show that strains lacking bL9 have fewer nuclease-resistant disomes, a hallmark of collisions, and cannot recruit SmrB as deeply into polysomes upon induction of collisions as the wild-type strain. These findings suggest that bL9 stabilizes collided ribosomes. Consistent with these findings, we show using stalling reporters that loss of bL9 has a small but reproducible negative impact on ribosome rescue pathways. To more broadly explore the role of bL9 in translation and the life of the cell, we revisited the antibiotic sensitivity and recoding phenotypes in the light of the bridging interaction observed in the disome structures. We use luciferase reporters based on the *ybeL* gene to show that the increases in recoding in bL9 mutant strains depend entirely on ribosome collisions. These studies reveal how bL9 acts to maintain translational reading frame and highlight the fact that ribosome collisions, while an important trigger of ribosome rescue pathways, pose a threat to translational fidelity.

## Results

### Mutations in bL9 at the collision interface sensitize cells to ribosome-targeting antibiotics

To study bL9’s function at the collision interface, we generated a knockout strain (ΔL9) of *E. coli* K12 MG1655 as well as several strains with point mutations designed to destabilize bL9’s interaction with the upstream ribosome. Mutation of the conserved Phe91 to Ala (F91A) is expected to reduce aromatic stacking interactions with 16S rRNA (Figure 1B) (15). In contrast, the conservative Phe91 to Tyr mutation (F91Y) should have a less destabilizing effect, especially given that some bacterial species have Tyr instead of Phe at this position (Figure S1), and the aromatic ring in Tyr may allow stacking to be maintained. Finally, we generated the Ser93 to Phe (S93F) mutant originally isolated in a genetic screen for increased recoding that is also found at the same interface (28). Each mutant allele was integrated into the genomic locus. The ΔL9 deletion and the three point mutations were all well-tolerated, with little or no change in the doubling times for the four mutant strains compared with wild-type in LB media (Figure 2A). In parallel, strains expressing FLAG-tagged versions of each bL9 variant were also created; an anti-FLAG blot confirmed that the bL9 mutants are expressed at levels comparable to the wild-type suggesting that the mutations are not destabilizing per se (Figure S2). The untagged bL9 variant strains were used for all subsequent assays, however, because the FLAG tag interferes with bL9 function (Figure S3).

**Figure 2.**
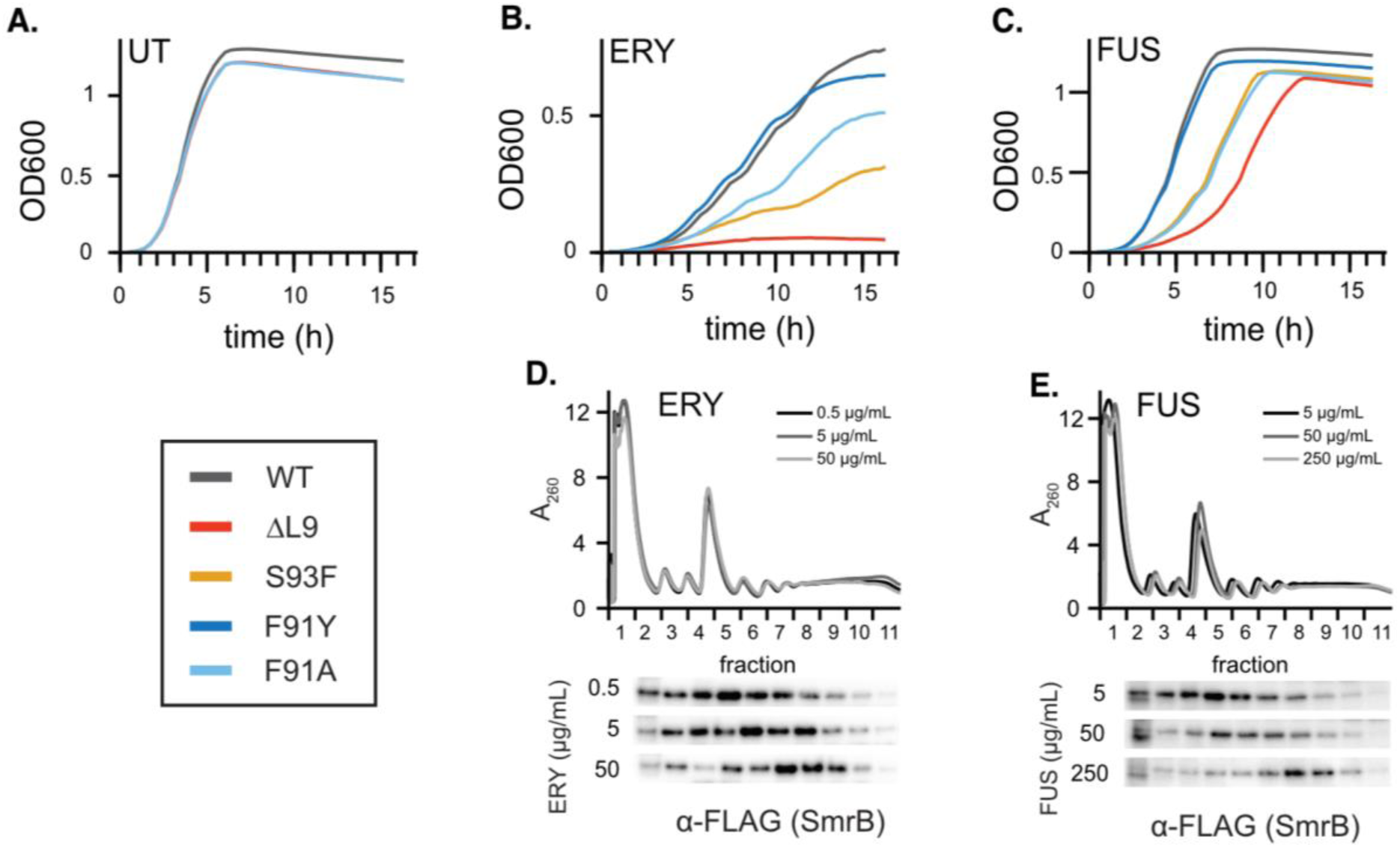
bL9 mutants are sensitive to antibiotics that cause ribosomal collisions. **(A-C)** Growth curves of bL9 mutants in various conditions: (A) untreated (UT), B) Erythromycin (ERY) 100 µg/mL, (C) Fusidic acid (FUS) 50 µg/mL. Curves are the mean of technical replicates from a representative experiment; the experiments were repeated at least three times for each drug. **(D, E)** FLAG-SmrB position within a sucrose gradient as an indicator of ribosome collisions. The WT bL9 FLAG-SmrB strain, bRZ118, was treated with (D) ERY or (E) FUS and lysates were separated on a sucrose gradient; polysome profiles are shown at the top of each panel. Western blots were performed on fractions from the gradients, probing with an α-FLAG antibody (Sigma) to detect FLAG-SmrB. The experiment was performed in triplicate; representative data from one replicate is shown.

We first characterized the effects of mutating bL9 in these strains by testing their sensitivity to antibiotics. Previous genome-wide studies reported that strains lacking bL9 are hypersensitive to ribosome-targeting antibiotics including macrolides and fusidic acid (FUS) (24, 25). We confirmed these findings, observing dramatic reductions in the growth rates for the ΔL9 strain in media containing the macrolide erythromycin (ERY) at 100 μg/mL (Figure 2B) or 50 μg/mL FUS (Figure 2C). The F91A and S93F mutants have intermediate phenotypes, showing a reduction in growth that is not as severe as the full deletion. In contrast, the sensitivity of the F91Y mutant strain is essentially indistinguishable from the wild-type strain, suggesting that this mutation is well tolerated, and the activity of bL9 is retained. The relative sensitivity of the bL9 variants to ERY and FUS is consistent across a range of drug concentrations (Figure S4). These results connect conserved residues at the collision interface with the antibiotic sensitivity observed when the bL9 is deleted.

We next showed that ERY and FUS induce ribosome collisions by following the distribution of SmrB in sucrose gradients after treating exponentially growing cultures with antibiotics for five minutes. Although SmrB can bind all ribosomes with low affinity, it is recruited deep into polysomes through specific interactions with both ribosomes at the collision interface (10). Previous studies have established that ERY induces robust levels of ribosome collisions through binding in the exit tunnel of the ribosome and blocking peptidyl transfer in a sequence dependent manner (10, 12, 39, 40). As expected, we find that at low ERY concentrations, well below the minimum inhibitory concentration (41), SmrB is distributed broadly across free subunits, 70S monosomes, and light polysomes (Figure 2D). But in higher concentrations of ERY, SmrB shifts deeper in the gradient and is enriched in the polysome fraction, providing strong evidence for ribosome collisions.

Given that FUS binds to EFG and traps it on the ribosome in a rotated state (27, 42), arresting elongation, we expected that FUS would also induce ribosome collisions. Indeed, we find that SmrB shifts from lighter fractions in sucrose gradients with sub-inhibitory levels of FUS to deeper fractions in the polysomes at higher FUS concentrations, consistent with high levels of ribosome collisions (Figure 2E). (Note that due to the mechanism of action of these drugs and their weak efficacy in *E. coli*, we cannot use very high concentrations to arrest all ribosomes in such a way that collisions cannot occur). We conclude that collisions are abundant at the chosen concentrations of both ERY and FUS. Taken together, these data establish that mutations at the collision interface reduce fitness in cells treated with antibiotics that induce ribosome collisions.

### bL9 stabilizes the interaction between collided ribosomes

One straightforward prediction of the structural models of collided disomes is that loss of the bL9 bridge between the two ribosomes should reduce their affinity for each other and therefore the stability of the disome pair. To observe collided disome levels, we treated cultures for 5 min with 5 µg/mL chloramphenicol (CAM), another elongation inhibitor which we previously showed induces high levels of ribosome collisions (12), then treated cell lysates with RNase to degrade unprotected mRNA, collapsing most of the polysomes to monosomes (Figure 3A). Collided disomes survive this treatment intact because the mRNA between them is protected from the nuclease by the collision interface (43, 44). As expected, lysates from wild-type *E. coli* show nuclease-resistant disomes and a smaller trisome peak (Figure 3A, right). In contrast, lysates from the ΔL9 knockout strain show a much-reduced nuclease-resistant disome peak and essentially no trisomes. These data are consistent with reduced stability of collided disomes in the ΔL9 knockout strain as previously reported (15). Lysates from the S93F mutant also show reduced abundance of nuclease-resistant disomes and trisomes, essentially indistinguishable from the ΔL9 knockout strain.

**Figure 3.**
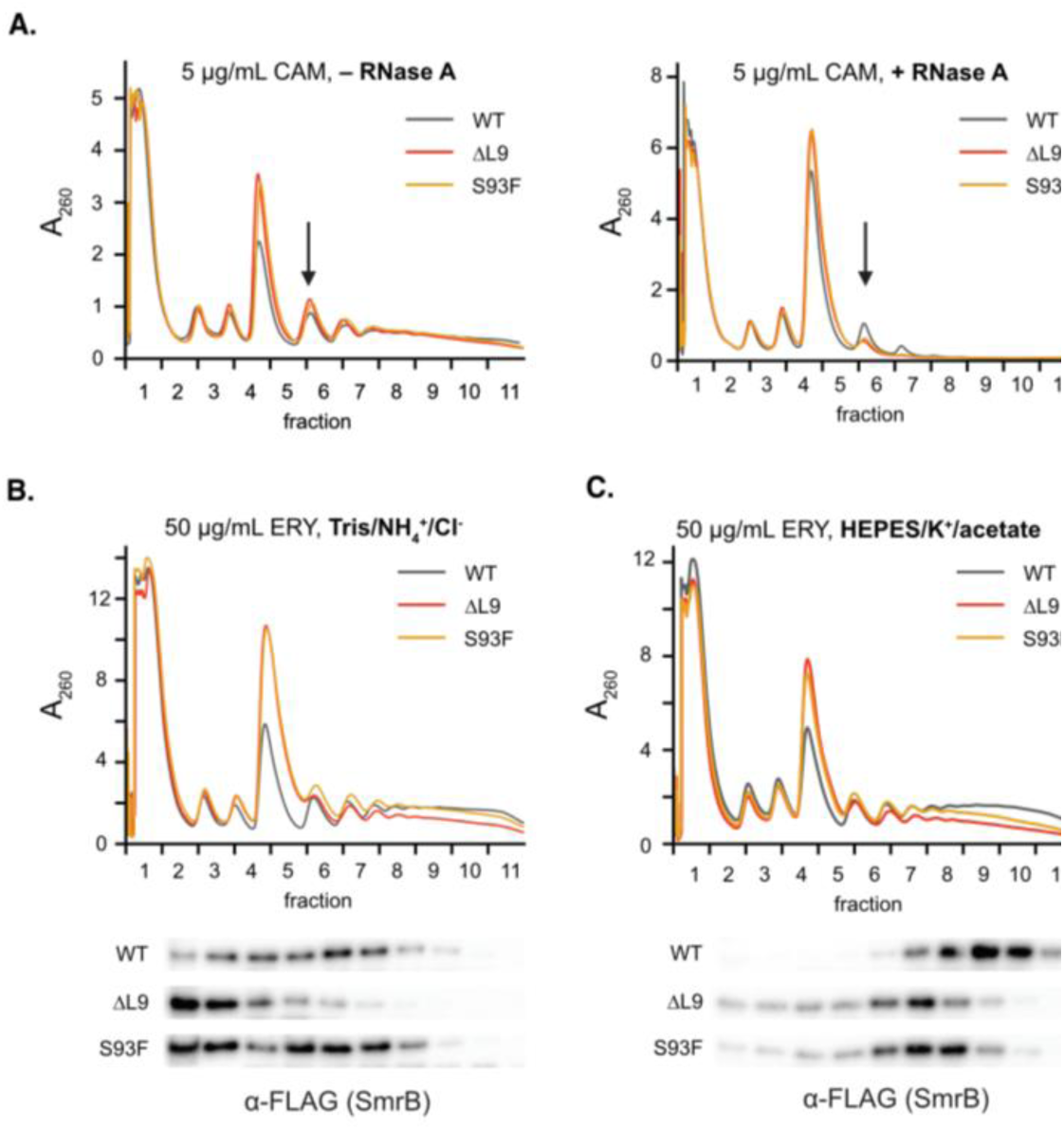
Loss of bL9 leads to unstable disomes. **(A)** Monitoring collisions via nuclease-resistant disomes. Cultures were treated with 5 µg/mL CAM for 5 minutes to induce ribosome collisions. Lysates were then left untreated (left) or treated with RNase A (right) and run on a sucrose gradient. Polysome traces are shown; the arrow indicates the disome peak. **(B, C)** FLAG-SmrB position within a sucrose gradient as an indicator of ribosome collisions. Cultures were treated with 50 µg/mL ERY to induce ribosome collisions, then lysates were run on a sucrose gradient to generate polysome profiles (top of each panel). Western blots were performed on fractions from the sucrose gradients, probing with an α-FLAG antibody (Sigma) to monitor FLAG-SmrB migration through the gradient. Standard lysis and gradient buffers containing Tris, NH4^+^ and MgCl_2_ were used in (B), whereas less stringent buffers containing HEPES, K^+^, and Mg(OAc)_2_ were used in (C). All experiments were performed in triplicate; representative data from one replicate is shown.

As a second approach for measuring levels of ribosome collisions, we asked how well SmrB is recruited to deep polysomes when these strains are treated with erythromycin (12). The expectation is that if the interface between the two collided ribosomes is less stable, SmrB will not be observed as deep in the sucrose gradient. Note that SmrB does not bind to bL9 directly in the cryo-EM structure of the complex (10). In wild-type cells treated with 50 µg/mL ERY, SmrB moves deep into polysomes, as expected (Figure 3B). Yet in the ΔL9 knockout strain, SmrB is found only in the light fractions, near the top of the gradient. The strain expressing the S93F mutant has an intermediate phenotype, with SmrB recruited less deeply than the wild-type, but more deeply than the knockout strain.

The dramatic loss of SmrB binding to ribosomes in the ΔL9 knockout strain led us to ask if our binding assay was too harsh; after all, there should be many ribosome collisions in ERY treated cells. Sucrose gradients are a demanding binding assay, as SmrB has ample opportunity to diffuse away from ribosomes moving through the gradient over the 2.5 h centrifugation. We therefore repeated the SmrB recruitment assay using sucrose gradients made with a less stringent buffer containing Mg(OAc)_2_ instead of MgCl_2_. In these gradients, SmrB shifts even deeper in the wild-type strain upon ERY treatment than in the gradients using the original buffer (Figure 3C). In the bL9 knockout and S93F strains, SmrB moves into the polysome fractions, indicating that these mutant ribosomes do not have gross SmrB binding defects. Yet once again, SmrB is present in somewhat lighter polysome fractions in the mutant strains than in the WT. Taken together, the reduced SmrB binding and reduced level of nuclease-resistant disomes observed in the mutant strains reveal that bL9 plays a role in stabilizing collided ribosomes.

### Strains lacking bL9 show a small reduction in ribosome rescue efficiency

We next asked if loss of bL9 affects the known ribosome rescue pathways. We reasoned that stabilization of the disome interface by bL9 might help ribosome rescue factors to recognize and clear the collision. To test this idea, we followed the activity of rescue factors SmrB and HrpA on a reporter where ribosomes stall at 12 rare Arg codons between the genes encoding *nanoLuc* and *ble* (Figure 4A). Previous work established that expression of the fusion protein from this stalling reporter is strongly inhibited by both SmrB, which cleaves the transcript leading to rapid mRNA decay, and HrpA, which splits stalled ribosomes into subunits, aborting translation (12). As expected, we found that a negative control with a reporter lacking a stalling sequence (Non-stall) shows little or no difference in expression in all four strains. With the Arg12 reporter, little full-length protein containing the C-terminal FLAG epitope is produced in the wild-type strain (Figure 4B) while in strains lacking either SmrB or HrpA, significantly higher levels of reporter protein accumulate because ribosomes are able to eventually translate through the Arg12 cluster and complete translation of the reporter (Figure 4B). Notably, there is also a small but reproducible increase in the full-length reporter levels in the ΔL9 knockout strain in comparison to the wild-type, consistent with a small reduction in ribosome rescue efficiency. The ability of SmrB to function in a ΔL9 strain is consistent with our earlier observation that FLAG-SmrB binds polysomes in the ΔL9 strain in the less stringent buffer. These results show that bL9 plays a modest but observable role in promoting efficient ribosome rescue, possibly by stabilizing the collision interface known to trigger ribosome rescue.

**Figure 4.**
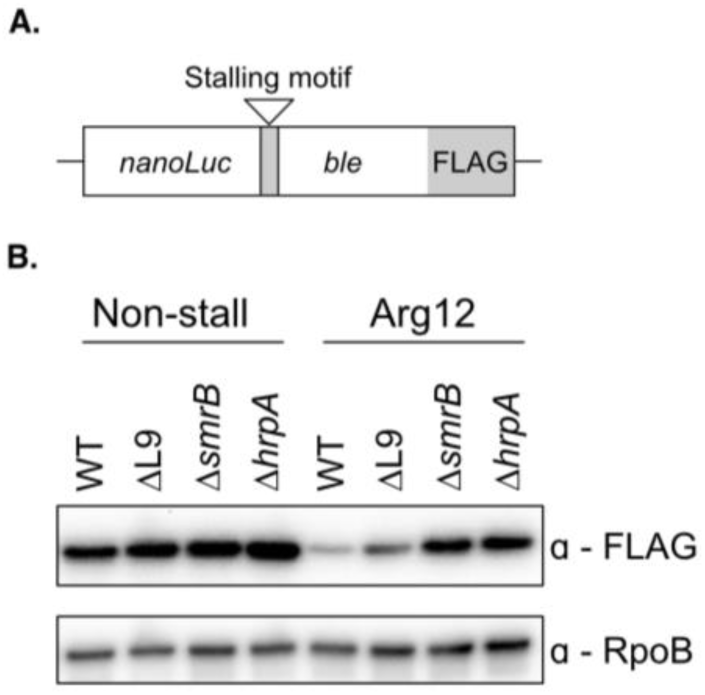
The effects of bL9 on ribosome rescue. **(A)** A schematic of reporter constructs containing a fusion of nanoLuc and FLAG proteins. The stalling reporter (pKS-Arg12) introduces a stalling motif with twelve rare Arg codons between the nanoLuc and Ble, whereas the Non-stall reporter (pKS-NonStall) has no additional sequence between them. **(B)** Western blot analysis showing the full-length reporter protein detected with an anti-FLAG antibody (Cell Signaling). RpoB serves as the loading control. All experiments were performed in triplicate; representative data from one replicate is shown.

### Mutations in bL9 at the collision interface affect recoding in gene 60

We also revisited the role of bL9 in suppressing recoding events in *gene 60* from T4 phage, the context where it was first shown to suppress recoding (28). We generated bypassing reporters which contain the full-length *gene 60* sequence containing all of the necessary components for bypassing (the nascent peptide signal, the 5’-stem loop, the take-off and landing sites, and the 3’-stem loop), fused to nanoluciferase such that luminescence is observed only upon successful recoding (Figure 5A). To validate the *gene 60*-*nanoLuc* reporter, we also constructed three reporters with mutations known to reduce bypassing efficiency, including the Stop codon mutant reporter (replacing the stop codon TAG with the sense codon TAC), the Take-off reporter (replacing the GGA take-off codon with GCA, creating a mismatch between the take-off and landing codons that can no longer be read by the same tRNA), and most importantly, the Stem-loop reporter which destabilizes the stem loop surrounding the take-off site (Figure 5B) (28, 30). High levels of luminescence were observed in wild-type cells expressing the wild-type *gene 60-nanoLuc* construct, with reduced signal from the control reporters with each of the mutations (Stop codon (8.1-fold), Take-off codon (7.4-fold), and Stem-loop (8.0-fold)) consistent with previous studies (Figure 5C) (28). These results demonstrate that luciferase activity from the reporter depends on bypassing in *gene 60*, as intended.

**Figure 5.**
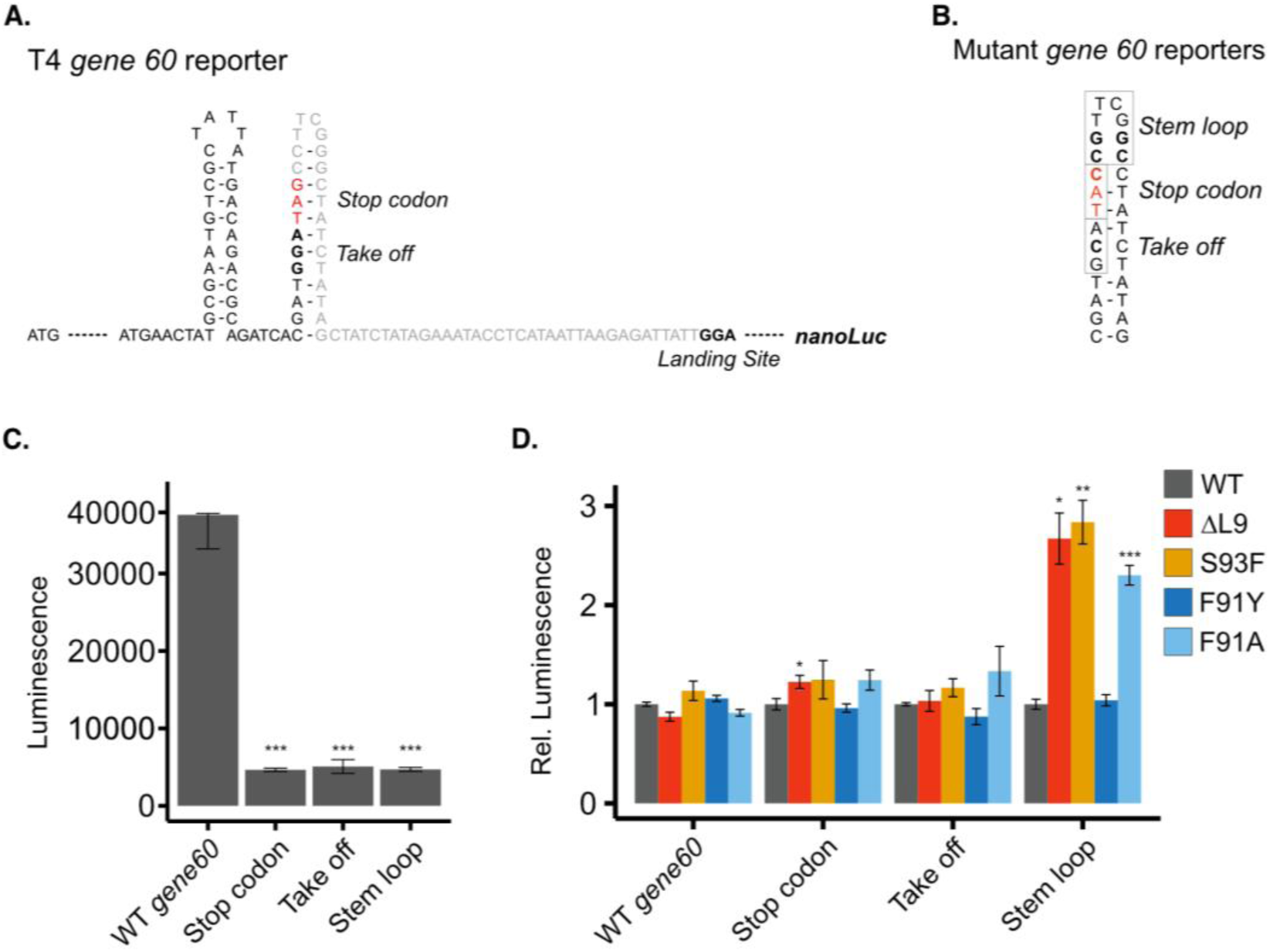
Recoding on *gene 60* increases in bL9 mutants. **(A)** The *gene60 – nanoLuc* reporter construct. **(B)** Boxes indicate location of the mutations in the *gene 60* sequence used for the Take off, Stop codon, and Stem loop reporters. **(C)** Luminescence of each of the *gene 60* reporters in the WT strain, bRZ137. **(D)** A panel of mutant gene 60 reporters was tested in various bL9 mutant strain backgrounds. The relative luminescence for each reporter is normalized to the signal of that reporter in the wild-type strain. Each bar reflects the mean ± the SE (C) or the SEM (D) of three biological replicates. (*) p < 0.05, (**) p < 0.01, (***) p < 0.001.

Using this series of reporters, we asked how defects in bL9 affect bypassing efficiency. To focus on comparisons between bL9 mutants, the data for each reporter are normalized to the wild-type strain. Deletion of bL9 has no significant effect on bypassing in the wild-type construct, where bypassing is highly efficient, nor in the Stop codon and Take-off controls, where loss of key mechanistic determinants leads to low levels of bypassing that cannot be restored by loss of bL9. But as expected, we see a 2.8-fold increase in bypassing efficiency in the Stem-loop reporter in the ΔL9 and S93F mutant strains (Figure 5D). This finding makes sense because the first evidence tying bL9 to recoding was obtained in a genetic screen for suppressors of the same Stem-loop mutant (28). The similarity of the phenotypes for the ΔL9 and S93F mutants suggests that S93F essentially leads to loss-of-function. Likewise, the F91A mutation leads to a 2.3-fold increase in bypassing on the Stem-loop reporter, while the F91Y mutation has no effect (Figure 5D). These data link bL9’s ability to suppress recoding to the region of the protein contacting the trailing ribosome in a collided disome.

### Recoding during termination in ybeL requires ribosome collisions

Testing whether the increase in recoding with bL9 mutations occurs in the context of ribosome collisions is difficult given the complex mechanism of *gene 60*. We therefore turned to the well-studied endogenous *E. coli* gene, *ybeL*, where ribosome stalling at the stop codon during termination leads to high levels of +1 frameshifting and hopping (35). We constructed a hopping reporter with the full-length *ybeL* gene (160 codons) where the C-terminal Pro is encoded by the CCG take-off codon. When the final codon is CCG, the ribosome can bypass or hop past the stop codon, resuming translation at another CCG codon 9 nt downstream (Figure 6A). We placed the *nanoLuc* gene downstream of the CCG landing codon so that luciferase activity reflects the efficiency of successful hopping. As a negative control, we created the CAG_hop reporter where the take-off codon is mutated from CCG to CAG. This mutation has two effects: it relieves stalling by replacing Pro with Gln, and reduces hopping, since the same tRNA cannot recognize CAG and CCG. As expected, when we normalize each reporter to the wild-type strain, we see little or no effect of the bL9 mutations on the CAG_hop control reporter. However, in the CCG_hop reporter there is a striking increase in bypassing in the ΔL9 strain (5.7-fold), and a smaller increase in bypassing in strains with the S93F, F91Y, or F91A mutations at the collision interface (2.0 to 2.5-fold) (Figure 6A). The increase in hopping or bypassing in bL9 mutants is consistent with our initial observations for the T4 *gene 60* reporter.

**Figure 6.**
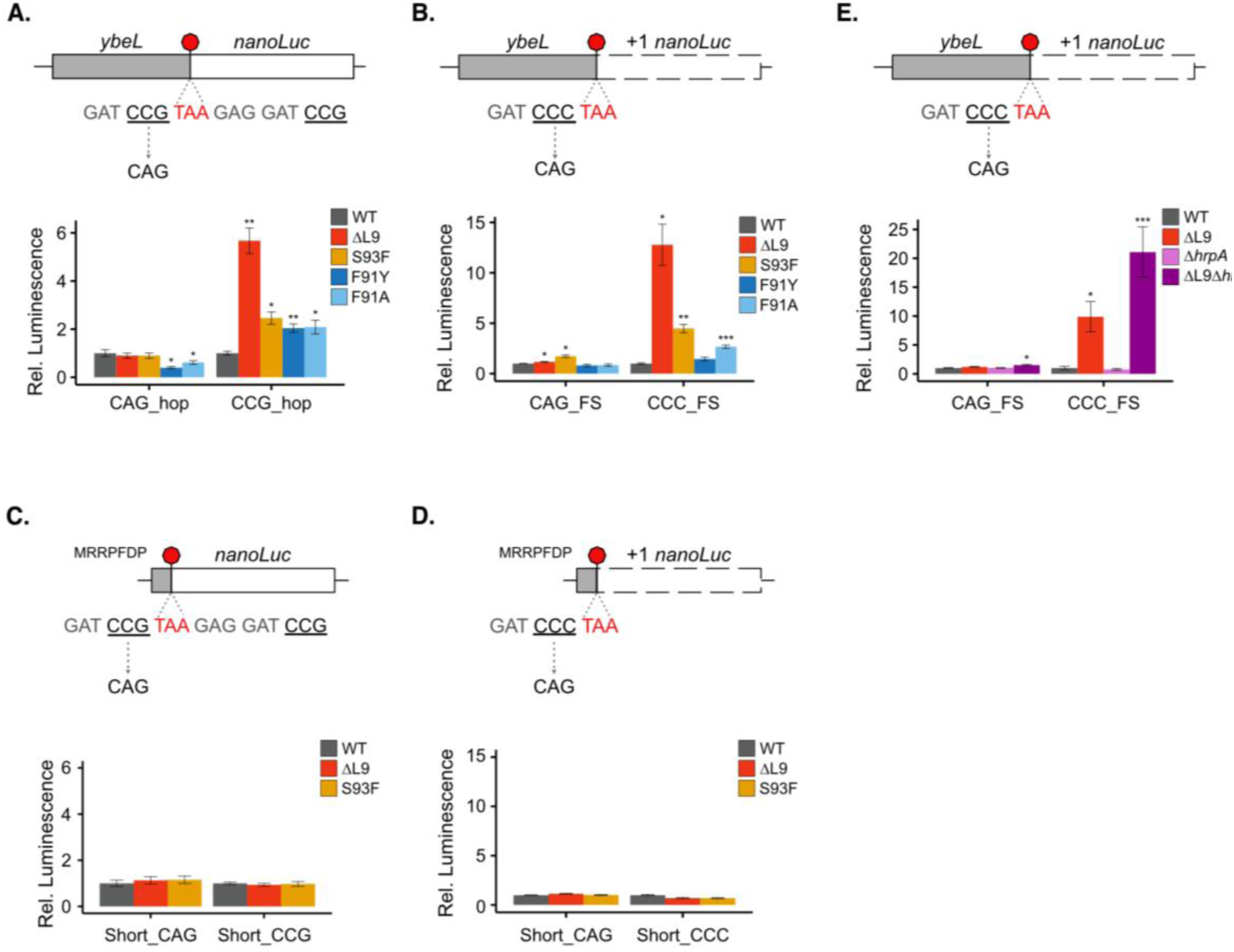
Recoding is collision and bL9 dependent. **(A,B)** *ybeL-nanoLuc* recoding reporters, used to monitor hopping or +1 frameshifting in bL9 mutant strains. **(A)** Hopping *ybeL – nanoLuc* reporters. The *ybeL* sequence ends in CCG-Stop (CCG_hop) followed by a matched landing CCG codon and the nanoLuc gene. The CAG_ hop serves as the negative control construct. **(B)** +1 frameshift (FS) reporter constructs where the *ybeL* gene ends in CCC-Stop (CCC_FS) with a downstream nanoLuc gene in the +1 frame. The CAG_FS serves as the negative control construct. **(C, D)** Short recoding reporters to monitor hopping or +1 frameshifting in bL9 mutant strains. The constructs encode a heptapeptide followed by a stop codon and then *nanoLuc*. Short_CCG (hop) or Short_CCC (FS) reporter constructs with nanoLuc downstream of the stop codon. The Short_CAG_hop or Short_CAG_FS constructs serve as negative controls. **(E)** The *ybeL*-*nanoLuc* +1 frameshift (FS) and control constructs used in panel B were introduced into strains with mutations in ribosome rescue factors. For each bar chart, the relative luminescence is normalized to the wild-type strain (bRZ137) for each reporter, and each bar reflects the mean ± the SEM of three biological replicates. (*) p < 0.05, (**) p <0.01, (***) p < 0.001.

We also created a reporter with the final Pro codon CCC allowing the ribosome to shift into the +1 frame at the end of *ybeL* and then translate *nanoLuc* (which is encoded in the +1 frame relative to *ybeL*) (Figure 6B). With the Pro codon in the P site and the stop codon in the A site during termination, prolonged stalling allows Pro-tRNA^GGG^, the P-site tRNA, to detach from the CCC codon, slide forward, and then pair with the +1 CCU codon, where the U comes from the first nt in the stop codon. After normalizing each reporter to the wild-type strain, little or no difference in the bL9 mutant strains is seen with the CAG_FS control. But with the CCC_FS reporter, there is 12.8-fold more frameshifting in the ΔL9 strain (Figure 6B). The point mutants at the collision interface also lead to higher levels of frameshifting, 4.5-fold for the original S93F mutant and 2.7-fold for the F91A mutant, with a minimal change for the conservative F91Y mutant (Figure 6B). Taken together, these data show that bL9 suppresses recoding in the content of both +1 frameshifting and hopping, consistent with the careful analyses of these outcomes upon ribosome stalling in *ybeL* as previously reported (35).

Unlike the complex bypassing mechanism in T4 *gene 60*, the fact that the stalling motif in our *ybeL* reporters is short (Asp-Pro-stop) allows us to ask whether collisions are required for the increases in bypassing and frameshifting in bL9 mutants. In previous studies, reporters with stalling motifs at various positions along the ORF were used to demonstrate that ribosome rescue factors only act on a stalled ribosome if the stalling motif is far enough from the start codon for multiple ribosomes to load and collide (8, 10, 12). Adapting the same strategy, we deleted all but the last seven residues of *ybeL* (MRRPFDP-stop) from the *ybeL*-*nanoLuc* reporters, so that there is not enough room to load a second ribosome upstream of the stalled ribosome (Figure 6C and D). Unlike the full-length constructs that allow for collisions, the short CCG_hop and CCC_FS reporters expressed in the wild-type, ΔL9, and S93F strains showed essentially no differences in recoding (Figure 6C and D). These data argue that the increase in bypassing and frameshifting seen in the bL9 mutants requires ribosome collisions.

If collisions promote recoding, then ribosome rescue factors that affect the level of collisions should also affect the level of recoding. Here, we focused on the helicase HrpA that splits stalled ribosomes into subunits; loss of HrpA leads to the accumulation of high levels of ribosome collisions (12). While no differences in luciferase activity were observed with the control reporter CAG_FS, we see a 2.1-fold increase in frameshifting on the full-length CCC_FS reporter in a ΔL9 Δ*hrpA* strain compared to the ΔL9 strain alone (Figure 6E). No increase in frameshifting was observed in the Δ*hrpA* strain, meaning that increased collisions do not lead to recoding at this site when bL9 is present and able to stabilize the collision interface (Figure 6E). As such, the ΔL9 deletion strain is sensitized to recoding defects in the context of collisions. The increase in frameshifting upon the loss of a ribosome rescue factor that resolves collisions further implicates collisions in recoding when bL9 is mutated.

### Reduced ribosome loading leads to fewer collisions and less recoding

An orthogonal approach for determining whether ribosome collisions increase recoding in bL9 mutants is to reduce ribosome loading on the reporters, such that stalling can be resolved before upstream ribosomes catch up and collisions occur (8, 45). We adopted this approach by mutating the Shine-Dalgarno sequence upstream of the start codon of the *ybeL*-*nanoLuc* reporters to vary initiation rates. At first, to identify ribosome binding sites with the desired levels of ribosome loading, we replaced the consensus Shine-Dalgarno motif AGGAGG with six random nucleotides upstream of the in-frame *ybeL-nanoLuc* reporter. After screening roughly 100 Shine-Dalgarno mutants for luciferase activity in the wild-type strain, we selected the following four for further study: the original consensus sequence High (AGGAGG, 100%), Medium (AGGACT, 82%), Low (GACGGT, 15%), and Very Low (CCTTTG, 1%) (Figure 7A). Note that these values reflect the difference in protein output due to lowered initiation rates for the in-frame *ybeL-nanoLuc* reporter, not differences in recoding.

**Figure 7.**
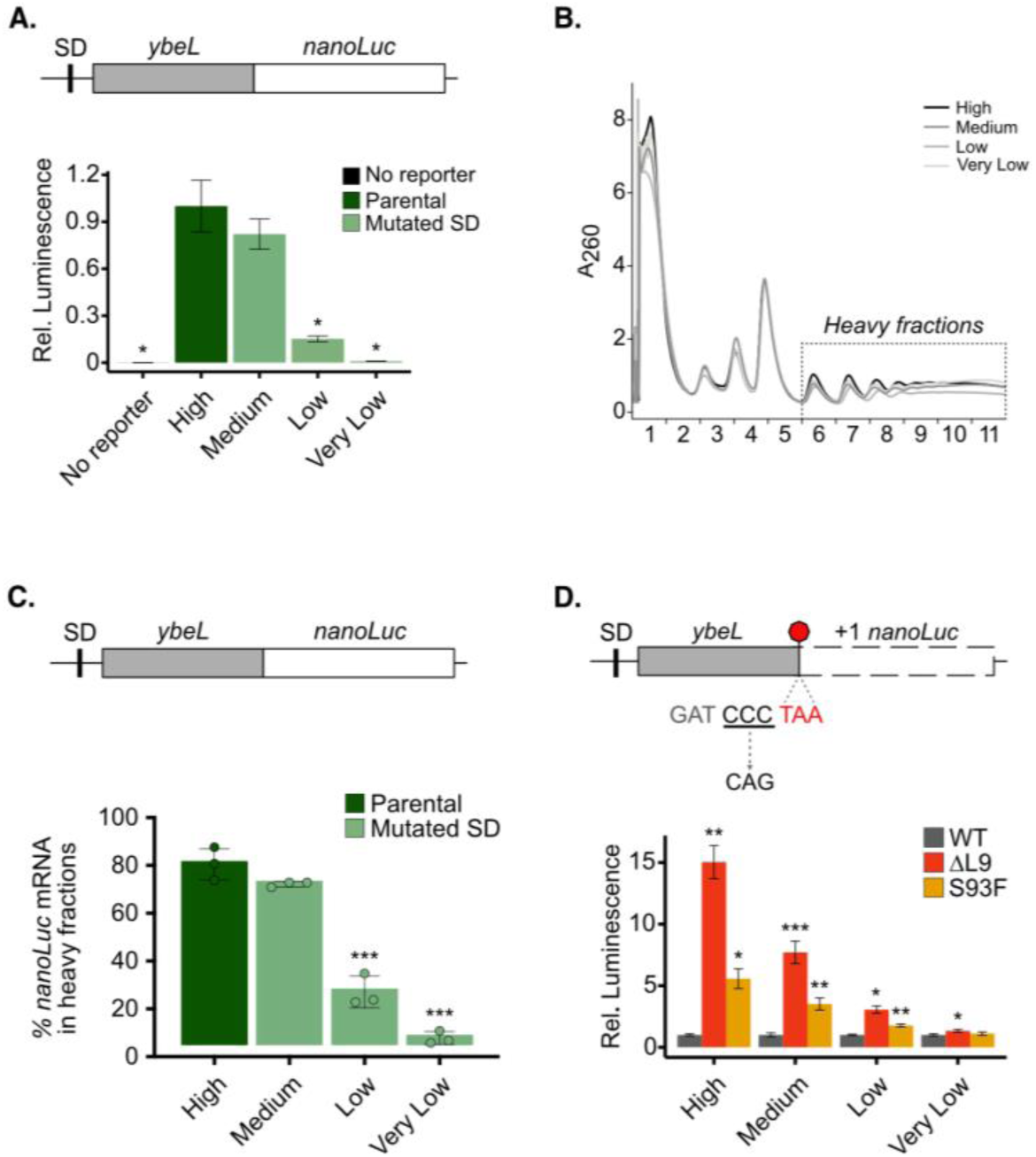
bL9 dependent recoding reduces with the ribosome load. **(A)** Shine Dalgarno (SD) reporters have varied SD sequences upstream of *ybeL – nanoLuc* in-frame fusion. The relative luminescence is normalized to the parental (AGGAGG) reporter; all reporters were expressed in DH5ɑ. **(B)** Representative polysome profiles of the SD reporters in DH5ɑ. Lysates for each SD reporter were run on a sucrose gradient. Profiles are left-aligned to the monosome peak. The disome and polysome fractions were pooled into the heavy fractions for **(C)** RT-qPCR across the gradient for the percentage of *nanoLuc* mRNA, normalized to an *eGFP* spike in. **(D)** +1 frameshift reporter constructs with varied SD sequences. The relative luminescence is normalized to the wild-type strain for each reporter (bRZ137). Each bar reflects the mean ± SEM of three biological replicates. (*) p < 0.05, (**) p < 0.01, (***) p < 0.001.

To further validate the translational efficiency of these reporters, we measured reporter mRNA levels in fractions from sucrose gradients (representative traces shown in Figure 7B). We performed RT-qPCR for the *nanoLuc* gene on RNA isolated from each fraction and normalized these values to an eGFP spike-in. We then calculated the percentage of reporter RNA found in the translating fractions (disomes and polysomes) (Figure S5). As expected, we see a decrease in the percentage of *nanoLuc* mRNA present within the heavy fractions in the reporters with weaker Shine-Dalgarno motifs (Figure 7C).

To ask how lowering initiation rates affects recoding, we next introduced these same Shine-Dalgarno sequences upstream of the *ybeL-nanoLuc* frameshifting reporter (CCC_FS) and measured luciferase activity in the wild-type, ΔL9, and S93F strains. As above, we see a nearly 15-fold increase in frameshifting in the ΔL9 strain with the same consensus SD motif used above (High, Figure 7D). However, the effects of bL9 on frameshifting gradually decrease as the initiation rate goes down, to 7.7-fold for the ΔL9 strain expressing the Medium reporter, to only 3.0-fold for the Low reporter, and essentially to no effect on the Very Low reporter (Figure 7D). The strong correlation between the initiation rates and ribosome loading from these four reporters with different Shine-Dalgarno sequences and the resulting recoding outcomes argue that lower ribosome loading (and fewer collisions) leads to lower frameshifting when bL9 is mutated. These data provide another line of support for the hypothesis that recoding occurs in the context of ribosome collisions.

## Discussion

Ribosome collisions are the signal to a cell that a problem has occurred (8, 46). The collision interface recruits rescue factors that trigger quality control pathways (7, 47). Ribosomes have evolved to stack together and remain tightly associated in roughly the same architecture in bacteria (10, 11), yeast (13), and mammalian cells (9). Here we explored the functional significance of one key difference at the interface in bacterial collided disomes: the bL9 bridge between the large subunit in the leading ribosome and the small subunit of the trailing ribosome. This bL9 bridge is observed in bacteria as different as *E. coli* and *B. subtilis* yet is not found in eukaryotes (10, 11). Our data suggest that this interaction contributes considerable stability to the collision interface, given that loss of this bridging interaction yields lower levels of nuclease-resistant disomes, as reported previously (15), and less recruitment of the collision sensor SmrB to deep polysomes. The lower stability of collided disomes in turn leads to somewhat lower ribosome rescue activity for SmrB or HrpA.

Rather than promoting ribosome rescue, we believe bL9’s main function at the collision interface is to suppress recoding events such as bypassing and frameshifting. Ribosomes lacking bL9 (or containing point mutants that disrupt its interactions with the trailing ribosome) show high levels of hopping and +1 frameshifting in our *ybeL-nanoLuc* reporters. Our recoding results are consistent with many earlier studies reporting that bL9 mutants have increased recoding at *gene 60* and *ybeL* (28, 35). The effect of bL9 on recoding has always been puzzling, especially given its distance from the active sites of the ribosome in a canonical monosome structure. Our work demonstrates that the increase is only observed in the context of ribosome collisions, and not at the start of the ORF (where there is no room to load a second ribosome) or when initiation rates are low (where a stalled ribosome resumes before a collision occurs).

While ribosome collisions are useful to the cell for highlighting problems in translation, these results reveal that paradoxically, collisions are also potentially deleterious. We envision that repeated attempts at translocation in the trailing ribosome may exert force on the leading ribosome, pulling the mRNA upstream or pushing the leading ribosome downstream (45). If this pulling force leads to loosening of the grip of the peptidyl-tRNA from the codon in the P site, the ribosome could slide downstream until base pairing is restored. This could explain the +1 frameshifting phenotype observed in the *ybeL-nanoLuc* reporters and the hopping events that require even more sliding along the mRNA. bL9 may prevent recoding in two ways (which are not mutually exclusive). First, it may further stabilize the disome interface, providing additional resistance against forces that destabilize the leading ribosome. This idea is consistent with our biochemical results showing that disomes are less stable when bL9 is mutated. Second, the bL9 binding to the trailing ribosome may sterically block the binding of EFG (19), preventing repeated attempts at translocation. This is also an appealing model, although we acknowledge that we do not have data to directly support it.

Our findings shed new light on old questions, including the decades-old observation that the S93F mutation in bL9 increases bypassing in the Stem loop mutant of *gene 60* (28). We recapitulated this increase for the Stem loop mutant in our reporter assays. Presumably the Stem loop mutant is defective specifically because the ribosome somehow struggles to slide down the mRNA, and this defect can be suppressed by ribosome collisions when the activity of bL9 is reduced or eliminated (32). In contrast, we did not observe increases in recoding for the wild-type *gene 60* construct or the other mutants, consistent with prior studies (28). In the context of wild-type *gene 60* bypassing is highly efficient, and it is unlikely that ribosome collisions play a role, given that the bypassing is highly over-determined, and no stalling at the stop codon is observed at the stop codon in single-molecule assays (33).

We believe that the hypersensitivity of strains lacking bL9 to fusidic acid may also be due to increased recoding triggered by collided ribosomes. Fusidic acid traps ribosomes in a chimeric state (CHI) late in translocation by preventing dissociation of EFG following GTP hydrolysis (42). In this chimeric state, the mRNA and tRNAs have translocated by one codon, but the small subunit is blocked from swiveling back into position to restore the classical (POST) state of the ribosome (27). We speculate that the CHI state trapped by fusidic acid is precarious and more susceptible to pulling forces in collided disomes than the canonical POST state, perhaps because contacts stabilizing peptidyl-RNA base paring in the P site are not fully formed. High levels of recoding on ribosomes trapped by FUS would lead to frameshifting and premature termination at stop codons, releasing incomplete nascent peptides with the potential for toxicity.

While bL9 forms a key interaction at the collision interface in bacteria, preventing recoding, eukaryotes have evolved a distinct solution, the protein known in yeast as Mbf1 (EDF1 in humans). Mbf1 mutants were isolated in a genetic screen for high levels of +1 frameshifting at stall sites in reporters in yeast (48). Several point mutations in uS3 (a protein in the mRNA entry channel) were also found to promote frameshifting. Later work revealed that these mutants only promote frameshifting in the context of ribosome collisions (45), using approaches similar to those used here. Mutations in uS3 prevent Mbf1 from binding to the ribosome, suggesting these genetic results are driven by the same basic underlying molecular mechanism (49).

Structural studies reveal that Mbf1 binds the trailing ribosome on the 40S subunit at the mRNA entry channel (50, 51), near the collision interface, with N-terminal residues contacting 40S in the leading ribosome (49). While the exact mechanism needs further characterization, one possibility is that Mbf1 binding to uS3 and helices in both the head and body of 40S locks the ribosome and prevents translocation (51). A conserved KKY motif and helical segment appear to clamp the mRNA, redirecting its path, perhaps improving the grip on the message (50). These phenotypes observed in yeast when Mbf1 is lost are remarkably similar to those observed in the absence of bL9 in *E. coli*, highlighting the fact that ribosome collisions are liable to induce recoding and can therefore be dangerous, and that suppressing translocation and mRNA slippage is necessary to maintain translational fidelity.

## Supplementary Data

Figures S1-S5

## Acknowledgments

*Author contributions:* Conceptualization, R.E.Z. and A.R.B.; Methodology & Investigation, R.E.Z. and J.L.J.; Writing – Original Draft, R.E.Z, J.L.J., and A.R.B.; Writing – Review & Editing, R.E.Z, J.L.J., A.R.B., and R.G.; Funding Acquisition, A.R.B. and R.G.; Supervision, A.R.B.

## Materials and Methods

### Reagents

All supplier and item information is available in Appendix Table 1. Biological Resources

### Strain growth and maintenance

*E. coli* strains were routinely grown at 37 °C in LB media, supplemented with 1.5% agar as needed for plates. Antibiotics were supplemented for strain maintenance at the following concentrations: carbenicillin (100 µg/mL) or kanamycin (50 µg/mL). Plasmids were routinely cloned into *E. coli* DH5alpha for storage.

### Plasmid and strain construction

Plasmids and primer sequences are included in Appendix Table 1 DNA knockouts and knock-in constructs were integrated into the *E. coli* genome using λ red recombinase (52); correct integration was verified by PCR across the locus and sequencing. Plasmids were introduced into chemically competent *E. coli* made using standard protocols (53) or the Zymo Mix & Go kit.

### Knockout of bL9

The ΔL9 strain was made by P1 transduction, moving the *rplI*::Kan^R^ locus from the *rplI* (L9) knockout strain in the Keio collection into K12 MG1655.

### Loopout of markers

To remove antibiotic resistance cassettes from *E. coli*, strains were transformed with pCP20, which encodes FLP recombinase, enabling recombination between the FRT sites which flank the resistance cassette. Strains were cultured in LB without drugs to allow for growth after loss of the resistance cassette, then grown at 42 °C to encourage loss of the pCP20 plasmid (54). Removal of the cassettes was verified by confirming appropriate drug sensitivities and by PCR (RZO1 & 2 for *rplI* locus) across the relevant genomic locus and sequencing the amplicons.

### FLAG-wtL9 plasmid construction

Initially, FLAG-tagged copy of wildtype *rplI* (bL9) was cloned into a plasmid, as a three-piece Gibson Assembly, creating pAC02. The backbone contains a p15ori and *amp^R^* marker, cut from pKS_Arg12 by restriction digest with SalI and HindIII. The *rplI* promoter and RBS were PCR-amplified from MG1655 genomic DNA, using primers pAC02/04 F1 and pACO2 R1. The *rplI* ORF was PCR-amplified using oligos pAC02 F2 and pACO2/4 R2. pAC02 R1 and pAC02 F2 are designed to introduce a FLAG tag at the N-terminus of the *rplI* gene.

### Cloning L9 variants

F91A or S93F point mutations were introduced into the FLAG-*rplI* gene in pAC02 using Quickchange Lightning kit, resulting in pRZ3 and pRZ6, respectively. pRZ3 was used as an inverse PCR template to create the FLAG-L9-F91Y plasmid, pRZ20. All plasmids were confirmed by sequencing.

A panel of plasmids lacking the FLAG tag but containing the same bL9 variants was also created. Quickchange lightning was used to remove the FLAG tag from pAC02 (oligos RZO 38 & 39), creating pRZ11. The plasmid was then digested XmaI/KpnI to isolate the N-terminal fragment of *rplI* (lacking the FLAG tag), and this was ligated into digested pRZ3, pRZ6, or pRZ20 to create untagged versions of the F91A, S93F, and F91Y variants of bL9 (pRZ14, pRZ18, pRZ22, respectively).

To integrate the bL9 variant alleles into the genome, we used fusion PCR to create a construct that contained the bL9 variant ORF followed by a KanR cassette. The bL9 variant was amplified from the corresponding plasmid, using a forward primer that maintained (RZO47) or lacked (RZO51) the FLAG tag at the N-terminus of *rplI*. The Kan^R^ cassette was amplified from pKD4 (RZO49/50) and includes the flanking FRT sites to allow for removal of the resistance cassette. These two fragments were then joined by a round of fusion PCR using RZO53 and RZO54. The knock-in construct was gel purified and transformed into a clean loopout ΔL9 strain, bRZ30c, using the lambda red system. Correct integration of the L9-KanR constructs was confirmed by PCR, and sequencing of the amplicon.

### Cloning nanoLuc reporters

The E_coli_STOP plasmid has an *ampR* marker, p15A origin of replication, and a BBa_J23107 promoter driving nanoLuc expression. Other nanoLuc reporter constructs were derived from this plasmid using Gibson assembly, inverse PCR, and the Quikchange Lightning Mutagenesis kit (Agilent). Further details of plasmid construction and the primer sequences used are in Appendix Table 1.

### Western Blots

Strains were grown to mid-log phase and pelleted by centrifugation. Cells were lysed by boiling in lysis buffer (12.5 mM Tris-Cl, pH 6.8, 4% SDS). The protein concentration of the resulting whole cell lysates were measured with a BCA assay, and samples were normalized prior to loading on a gel. Samples were run on 4-12 % BisTris SDS-PAGE midi gels, in MES buffer at 150 V for ∼ 50 min, with Precision Plus Protein Dual Color standards. Samples were transferred to PVDF membranes in a Transblot Turbo system, using the low MW program. Membranes were blocked in TBST + 5 % milk. Antibodies used were α-FLAG-HRP (Sigma or Cell Signaling), α-RpoB (BioLegend), and goat-α-mouse-HRP (Invitrogen) was used as a secondary when needed. Signal was detected using Thermo SuperSignal Pico Chemiluminescent substrate, supplemented with Femto as needed. Images were captured on a BioRad ChemiDoc imager.

### Growth assays

Single colonies of the desired strain were inoculated into 2 mL LB (for unmarked strains) or LB+Kan (for knockout or knock-in strains) and grown at 37 °C for ∼ 3 h so they were actively growing. In a 96-well plate, LB media supplemented with the desired drug titrated to various concentrations was dispensed across the plate, with at least two replicate wells per drug per strain. Actively growing cells were inoculated at a starting OD_600_ = 0.01, in a final volume of 150 µL per well. The 96-well plate was lidded, and growth was monitored in a BioTek Synergy H1 microplate reader for 16 h at 37 °C, with linear shaking. OD_600_ readings were taken every 20 min. The resulting data was analyzed in Python: The average OD reading for all blank wells was subtracted from the readings of the inoculated wells. The OD of technical duplicate wells were averaged and plotted over time for each strain.

### Polysome profiling

*E. coli* strains were grown to mid-log phase in LB media at 37 °C shaking. The culture was treated with the indicated drug for 5 min, at concentrations indicated in the text and figures. Cells were filter harvested as described previously (12), flash frozen in liquid N_2_ and stored at −80 °C. Cells were lysed in gradient lysis buffer (20 mM Tris pH 8, 10 mM MgCl_2_, 100 mM NH_4_Cl, 5 mM CaCl_2_, 0.4% TritonX 100, 0.1% NP40, DNAse I (10 U/mL), CAM (0.34 mg/mL) using a freezer mill, and resulting lysates were clarified by centrifugation at 18210 x g at 4 °C for 10 min. A_260_ measurements were taken with a Nanodrop and used to normalize the lysates between strains. A 10-50% sucrose gradient was made in standard gradient buffer (20 mM Tris pH 8, 10 mM MgCl_2_, 100 mM NH_4_Cl, 2 mM DTT) using a Gradient Maker. 35 A.U. of lysate were loaded onto each sucrose gradient, and samples were centrifuged in an SW-41 rotor at 35,000 rpm at 4° C for 2.5 h. The gradients were fractionated using a Piston fractionator (BioComp), which records A_260_ readings during collection. For experiments where the fractions are processed for Western blots, trichloroacetic acid (TCA) was added to 10% final at the time of fraction collection. TCA precipitation and Western blotting followed, to track FLAG-SmrB across the gradient (see below).

### Modified lysis & gradients

For Figure 3c, cultures were grown and samples were processed as described above in Polysome Profiling but instead used modified lysis and gradient buffers containing HEPES, K^+^, and acetate, instead of our standard buffer recipes. The HEPES lysis buffer contained 20 mM HEPES pH 8, 10 mM Magnesium acetate, 100 mM potassium acetate, 5 mM calcium acetate, 0.4% TritonX 100, 0.1% NP40, DNAse I (10 U/mL), CAM (0.34 mg/mL). The HEPES gradient buffer contained 20 mM HEPES pH 8, 10 mM Magnesium acetate, 100 mM potassium acetate, 2 mM DTT.

### TCA Precipitation

Proteins were concentrated from fractionated gradients. 1 mL fractions were collected with 10% TCA (See polysome profiling) and precipitated overnight at −20 °C. Samples were centrifuged at 18213 x g for 10 min at 4 °C and the buffered sucrose was removed from the pellet by aspiration. The pellets were washed twice in 500 µL ice-cold acetone and spun as before. The protein pellet was allowed to air dry at RT (∼ 5 min) and then resuspended in 30 µL SBTU (2x SB, 100 mM Tris pH 8.0, 6 M urea, by heating at 37 °C for 30 min, shaking at 1200 rpm. 5 µL of the precipitated fractions 2-12 were loaded onto a gel for SDS-PAGE and Western blotting (see above). Fraction 1 was omitted from the gels due to a large quantity of non-specific signal from the anti-FLAG antibodies on the Western blot, which interfered with detection of the FLAG-SmrB signal in other fractions from the gradient. Note the samples with urea in the buffer were not heated prior to loading on the SDS-PAGE gel.

### Nuclease resistant disomes

Strains were grown in 100 mL LB to mid-log, treated for 5 minutes with 5 µg/mL chloramphenicol to induce collisions, and harvested by filtration. Cells were lysed in standard lysis buffer, and sample concentrations were normalized based on A_260_ readings. For each sample, two 10 A.U. aliquots of each sample were taken. One was saved as an untreated (-RNase A) sample, and the other was treated with 0.1x volume of RNase A (ThermoFisher) (diluted to 10 µg/ml in lysis buffer) at 25 °C shaking at 900 rpm for 1 h. RNase digestion was stopped by adding 0.01x volume of SuperaseIn nuclease inhibitor (Invitrogen). 10 A.U. of the sample (± RNase A) were loaded onto 10-50% sucrose gradients in standard gradient buffer and centrifuged as described in “Polysome Profiling” above. After centrifugation, the piston fractionator was used to read A_260_ readings across the gradient, but the fractions were not collected.

### Luminescence assays

Cultures were grown to mid-log phase and normalized by OD600 before mixing 45 µL of each culture with 5 µL of a high salt buffer (1 M K_2_HPO_4_ pH 7.8, 20 mM EDTA). Samples were lysis via freeze thaw on dry ice and then mixed with 150 µL of lysis buffer (1X Luciferase Cell Culture Lysis Reagent (Promega), 1.25 mg/mL lysozyme, 2.5 mg/mL BSA). Equal volumes of sample and luciferase reagent were prepared according to manufacturer’s instructions using the Nano-Glo® Luciferase Assay System (Promega). The plate was loaded into the BioTek Synergy H1 microplate reader, shaken for 3 min at 800 cpm, then after 7 min of settling, luminescence was measured. The resulting data was analyzed in R. Absorbance readings for blank wells were subtracted from the sample wells; relative luminescence between strain backgrounds was normalized to the wild-type strain for each reporter. Welch’s t-tests were used to calculate p-values.

### RNA extractions and RT-qPCR

RNA was purified from standard MgCl_2_ sucrose gradients by adding 500 µL of each fraction to 1 mL of TRIZOL (Invitrogen) containing 50 pg of CleanCap® EGFP mRNA spike-in (TriLink). Samples were frozen at −80 °C, thawed, and separated by chloroform and centrifugation. 500 µL of the aqueous layer was precipitated with isopropanol, and washed with 75% ethanol, and resuspended in water. 2 µg of the RNA was used in the reverse transcription which was performed according to the manufacturers of the iScript™ cDNA Synthesis Kit (BioRad), then diluted 3-fold in water.

RT-qPCR samples were performed by assembling 4.3 µL of the diluted cDNA product, 0.7 µL of the primer set (final primer concentration of 350 nM), and 5 µL of iTaq Universal SYBR Green Supermix (BioRad), then performed using the QuantStudio 6 Flex System with a protocol of: 95 °C for 2:20 min, 40 x (95 °C for 10 s, 60 °C for 30 s), 95 °C for 15 s, 60 °C for 1 min, to 95 °C at 0.05 °C/s for 15 s. For each primer set a standard curve was made by using dilute SD_AGGAGG plasmid or EGFP-pBAD (EGFP-pBAD was a gift from Michael Davidson (Addgene plasmid # 54762). Primer sequences are provided in Appendix Table 1. Using sPython, the relative amount of *nanoLuc* mRNA in each fraction was normalized to the EGFP spike in, and the percentage of the *nanoLuc* mRNA in each fraction pool (translating or non-translating) was determined. P-values were calculated using Welch’s t-tests.

## Statistical Analyses

Growth curves of L9 mutants in FUS or ERY were performed in triplicate; one representative experiment is shown, with the curve representing the average between technical replicate wells. Figure 3 experiments, including polysome profiles on L9 variants were completed in triplicate, and one representative experiment is shown. Western blots detecting FLAG-SmrB across sucrose gradients were performed in at least triplicate.

## References

1. Yan, L.L. and Zaher, H.S. (2019) How do cells cope with RNA damage and its consequences? J. Biol. Chem., 294, 15158–15171.

2. Li, X., Hirano, R., Tagami, H., et al. (2006) Protein tagging at rare codons is caused by tmRNA action at the 3′ end of nonstop mRNA generated in response to ribosome stalling. RNA, 12, 248–255.

3. Roche, E.D. and Sauer, R.T. (1999) SsrA-mediated peptide tagging caused by rare codons and tRNA scarcity. EMBO J., 18, 4579–4589.

4. Ito, K. and Chiba, S. (2013) Arrest Peptides: *Cis*-Acting Modulators of Translation. Annu. Rev. Biochem., 82, 171–202.

5. Wilson, D.N. (2014) Ribosome-targeting antibiotics and mechanisms of bacterial resistance. Nat. Rev. Microbiol., 12, 35–48.

6. Lin, J., Zhou, D., Steitz, T.A., et al. (2018) Ribosome-Targeting Antibiotics: Modes of Action, Mechanisms of Resistance, and Implications for Drug Design. Annu. Rev. Biochem., 87, 451–478.

7. Buskirk, A.R. (2026) Ribosome Collisions Trigger Ribosome Rescue in Bacteria. Annu. Rev. Microbiol., 80.

8. Simms, C.L., Yan, L.L. and Zaher, H.S. (2017) Ribosome Collision Is Critical for Quality Control during No-Go Decay. Mol. Cell, 68, 361–373.e5.

9. Juszkiewicz, S., Chandrasekaran, V., Lin, Z., et al. (2018) ZNF598 Is a Quality Control Sensor of Collided Ribosomes. Mol. Cell, 72, 469–481.e7.

10. Saito, K., Kratzat, H., Campbell, A., et al. (2022) Ribosome collisions induce mRNA cleavage and ribosome rescue in bacteria. Nature, 603, 503–508.

11. Cerullo, F., Filbeck, S., Patil, P.R., et al. (2022) Bacterial ribosome collision sensing by a MutS DNA repair ATPase paralogue. Nature, 603, 509–514.

12. Campbell, A., Esser, H.F., Burroughs, A.M., et al. (2025) The RNA helicase HrpA rescues collided ribosomes in E. coli. Mol. Cell, 85, 999–1007.e7.

13. Ikeuchi, K., Tesina, P., Matsuo, Y., et al. (2019) Collided ribosomes form a unique structural interface to induce Hel2-driven quality control pathways. EMBO J., 38, EMBJ2018100276.

14. Park, E.N., Mackens-Kiani, T., Berhane, R., et al. (2024) B. subtilis MutS2 splits stalled ribosomes into subunits without mRNA cleavage. EMBO J., 43, 484–506.

15. Flügel, T., Schacherl, M., Unbehaun, A., et al. (2024) Transient disome complex formation in native polysomes during ongoing protein synthesis captured by cryo-EM. Nat. Commun., 15, 1756.

16. Xue, L., Lenz, S., Zimmermann-Kogadeeva, M., et al. (2022) Visualizing translation dynamics at atomic detail inside a bacterial cell. Nature, 610, 205–211.

17. Xue, L., Spahn, C.M.T., Schacherl, M., et al. (2025) Structural insights into context-dependent inhibitory mechanisms of chloramphenicol in cells. Nat. Struct. Mol. Biol., 32, 257–267.

18. Schuwirth, B.S., Borovinskaya, M.A., Hau, C.W., et al. (2005) Structures of the Bacterial Ribosome at 3.5 Å Resolution. Science, 310, 827–834.

19. Selmer, M., Gao, Y.-G., Weixlbaumer, A., et al. (2012) Ribosome engineering to promote new crystal forms. Acta Crystallogr. D Biol. Crystallogr., 68, 578–583.

20. Yusupov, M.M., Yusupova, G.Zh., Baucom, A., et al. (2001) Crystal Structure of the Ribosome at 5.5 Å Resolution. Science, 292, 883–896.

21. Fischer, N., Neumann, P., Konevega, A.L., et al. (2015) Structure of the E. coli ribosome–EF-Tu complex at <3 Å resolution by Cs-corrected cryo-EM. Nature, 520, 567–570.

22. Baba, T., Ara, T., Hasegawa, M., et al. (2006) Construction of Escherichia coli K-12 in-frame, single-gene knockout mutants: the Keio collection. Mol. Syst. Biol., 2, MSB4100050.

23. Liu, A., Tran, L., Becket, E., et al. (2010) Antibiotic Sensitivity Profiles Determined with an Escherichia coli Gene Knockout Collection: Generating an Antibiotic Bar Code. Antimicrob. Agents Chemother., 54, 1393–1403.

24. Nichols, R.J., Sen, S., Choo, Y.J., et al. (2011) Phenotypic Landscape of a Bacterial Cell. Cell, 144, 143–156.

25. Ma, Y., Pirolo, M., Jana, B., et al. (2024) The intrinsic macrolide resistome of *Escherichia coli*. Antimicrob. Agents Chemother., 68, e00452–24.

26. Gao, Y.-G., Selmer, M., Dunham, C.M., et al. (2009) The Structure of the Ribosome with Elongation Factor G Trapped in the Posttranslocational State. Science, 326, 694–699.

27. Zhou, J., Lancaster, L., Donohue, J.P., et al. (2013) Crystal Structures of EF-G–Ribosome Complexes Trapped in Intermediate States of Translocation. Science, 340, 1236086.

28. Herbst, K.L., Nichols, L.M., Gesteland, R.F., et al. (1994) A mutation in ribosomal protein L9 affects ribosomal hopping during translation of gene 60 from bacteriophage T4. Proc. Natl. Acad. Sci., 91, 12525–12529.

29. Huang, W.M., Ao, S.-Z., Casjens, S., et al. (1988) A Persistent Untranslated Sequence Within Bacteriophage T4 DNA Topoisomerase Gene 60. Science, 239, 1005–1012.

30. Weiss, R.B., Huang, W.M. and Dunn, D.M. (1990) A nascent peptide is required for ribosomal bypass of the coding gap in bacteriophage T4 gene *60*. Cell, 62, 117–126.

31. Herr, A.J., Gesteland, R.F. and Atkins, J.F. (2000) One protein from two open reading frames: mechanism of a 50 nt translational bypass. EMBO J., 19, 2671–2680.

32. Herr, A.J., Nelson, C.C., Wills, N.M., et al. (2001) Analysis of the roles of tRNA structure, ribosomal protein L9, and the bacteriophage T4 gene 60 bypassing signals during ribosome slippage on mRNA. J. Mol. Biol., 309, 1029–1048.

33. Samatova, E., Konevega, A.L., Wills, N.M., et al. (2014) High-efficiency translational bypassing of non-coding nucleotides specified by mRNA structure and nascent peptide. Nat. Commun., 5, 4459.

34. Chen, J., Coakley, A., O’Connor, M., et al. (2015) Coupling of mRNA Structure Rearrangement to Ribosome Movement during Bypassing of Non-coding Regions. Cell, 163, 1267–1280.

35. Seidman, J.S., Janssen, B.D. and Hayes, C.S. (2011) Alternative Fates of Paused Ribosomes during Translation Termination. J. Biol. Chem., 286, 31105–31112.

36. Hayes, C.S., Bose, B. and Sauer, R.T. (2002) Proline Residues at the C Terminus of Nascent Chains Induce SsrA Tagging during Translation Termination. J. Biol. Chem., 277, 33825–33832.

37. Woolstenhulme, C.J., Parajuli, S., Healey, D.W., et al. (2013) Nascent peptides that block protein synthesis in bacteria. Proc. Natl. Acad. Sci., 110, E878–E887.

38. Smith, A.M., Costello, M.S., Kettring, A.H., et al. (2019) Ribosome collisions alter frameshifting at translational reprogramming motifs in bacterial mRNAs. Proc. Natl. Acad. Sci., 116, 21769–21779.

39. Kannan, K., Kanabar, P., Schryer, D., et al. (2014) The general mode of translation inhibition by macrolide antibiotics. Proc. Natl. Acad. Sci., 111, 15958–15963.

40. Beckert, B., Leroy, E.C., Sothiselvam, S., et al. (2021) Structural and mechanistic basis for translation inhibition by macrolide and ketolide antibiotics. Nat. Commun., 12, 4466.

41. Li, X., Gan, Q. and Fan, C. (2026) Genome-wide screening of antibiotic survival genes in Escherichia coli MG1655. BMC Microbiol., 26, 152.

42. Belardinelli, R. and Rodnina, M.V. (2017) Effect of Fusidic Acid on the Kinetics of Molecular Motions During EF-G-Induced Translocation on the Ribosome. Sci. Rep., 7, 10536.

43. Wolin, S.L. and Walter, P. (1988) Ribosome pausing and stacking during translation of a eukaryotic mRNA. EMBO J., 7, 3559–3569.

44. Guydosh, N.R. and Green, R. (2014) Dom34 Rescues Ribosomes in 3′ Untranslated Regions. Cell, 156, 950–962.

45. Simms, C.L., Yan, L.L., Qiu, J.K., et al. (2019) Ribosome Collisions Result in +1 Frameshifting in the Absence of No-Go Decay. Cell Rep., 28, 1679–1689.e4.

46. Ferrin, M.A. and Subramaniam, A.R. (2017) Kinetic modeling predicts a stimulatory role for ribosome collisions at elongation stall sites in bacteria. eLife, 6, e23629.

47. Inada, T. and Beckmann, R. (2024) Mechanisms of Translation-coupled Quality Control. J. Mol. Biol., 436, 168496.

48. Wang, J., Zhou, J., Yang, Q., et al. (2018) Multi-protein bridging factor 1(Mbf1), Rps3 and Asc1 prevent stalled ribosomes from frameshifting. eLife, 7, e39637.

49. Kim, K.Q., Li, J.J., Nanjaraj Urs, A.N., et al. (2024) Multiprotein bridging factor 1 is required for robust activation of the integrated stress response on collided ribosomes. Mol. Cell, 84, 4594–4611.e9.

50. Sinha, N.K., Ordureau, A., Best, K., et al. (2020) EDF1 coordinates cellular responses to ribosome collisions. eLife, 9, e58828.

51. Pochopien, A.A., Beckert, B., Kasvandik, S., et al. (2021) Structure of Gcn1 bound to stalled and colliding 80S ribosomes. Proc. Natl. Acad. Sci., 118, e2022756118.

52. Datsenko, K.A. and Wanner, B.L. (2000) One-step inactivation of chromosomal genes in Escherichia coli K-12 using PCR products. Proc. Natl. Acad. Sci., 97, 6640–6645.

53. Green, R. and Rogers, E.J. (2013) Transformation of Chemically Competent E. coli. In Lorsch,J. (ed), Laboratory Methods in Enzymology: DNA, Methods in Enzymology. Academic Press, Vol. 529, pp. 329–336.

54. Cherepanov, P.P. and Wackernagel, W. (1995) Gene disruption in *Escherichia coli*: TcR and KmR cassettes with the option of Flp-catalyzed excision of the antibiotic-resistance determinant. Gene, 158, 9–14.

