## Supplemental Figures for "bL9 prevents recoding in collided ribosomes"

### Table of Contents:

Figure S1. Protein sequence alignment of bL9 page 1

Figure S2. FLAG-bL9 expression levels are consistent page 2

Figure S3. FLAG-tagged bL9 has a growth defect page 3

Figure S4. bL9 mutant growth curves page 4

Figure S5. Distribution of *nanoLuc* mRNA within each fraction. page 5

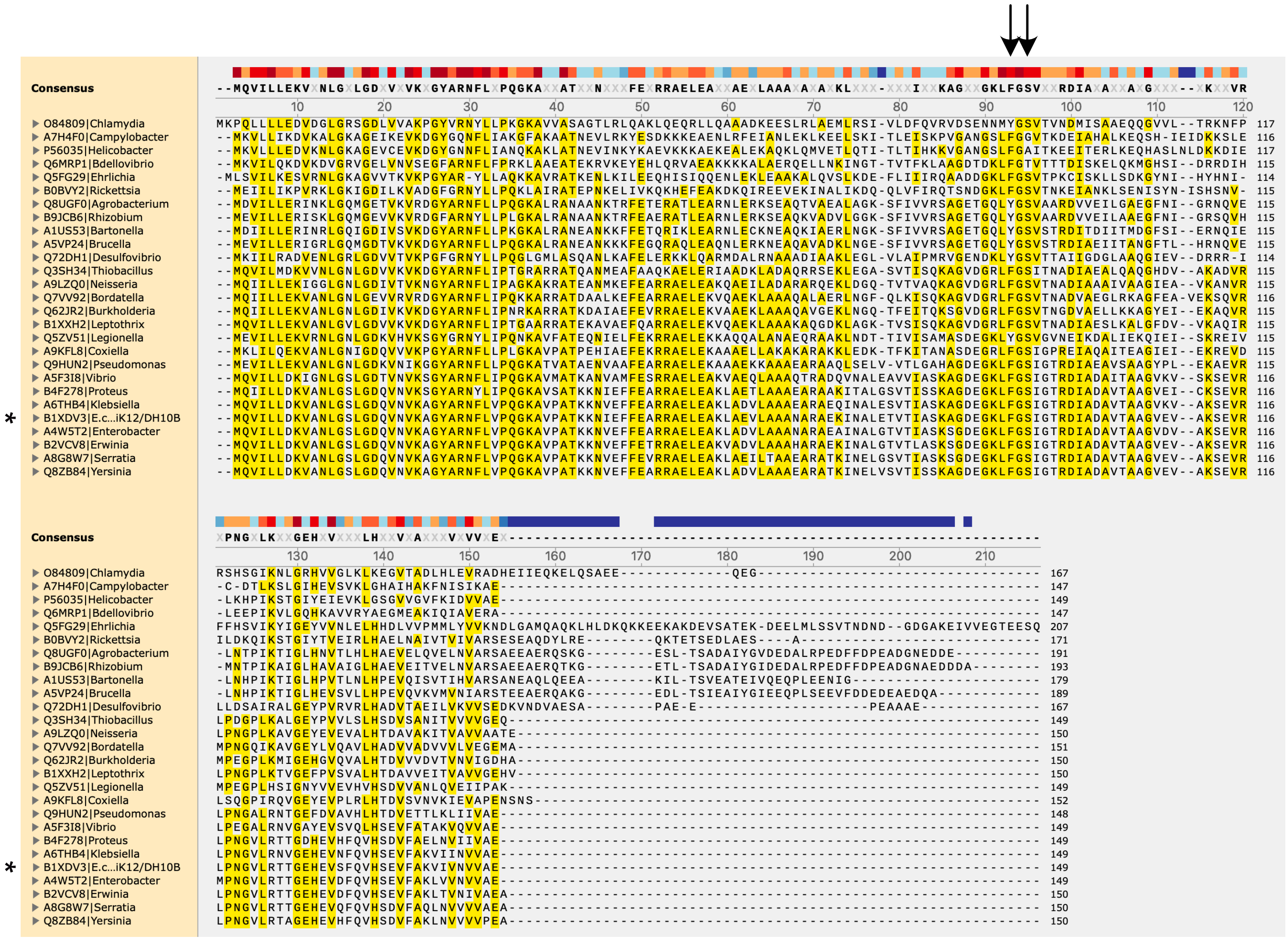


Figure S1. Protein sequence alignment of bL9 from Pseudomonadota (proteobacteria).

Interpro protein sequences from family "large ribosomal subunit protein bL9 (IPR000244)" were downloaded and twenty seven species were chosen from different genera, with representatives across alpha, beta, gamma, delta, and epsilon proteobacteria. Protein sequences were aligned with Clustal-Omega on EBI website (https://www.ebi.ac.uk/jdispatcher/msa/clustalo). The consensus threshold was set to 50%, and yellow highlighting denotes residues that match the consensus.

* *E. coli* protein sequence. Arrows mark the residues F91 and S93 from the *E. coli* sequence.

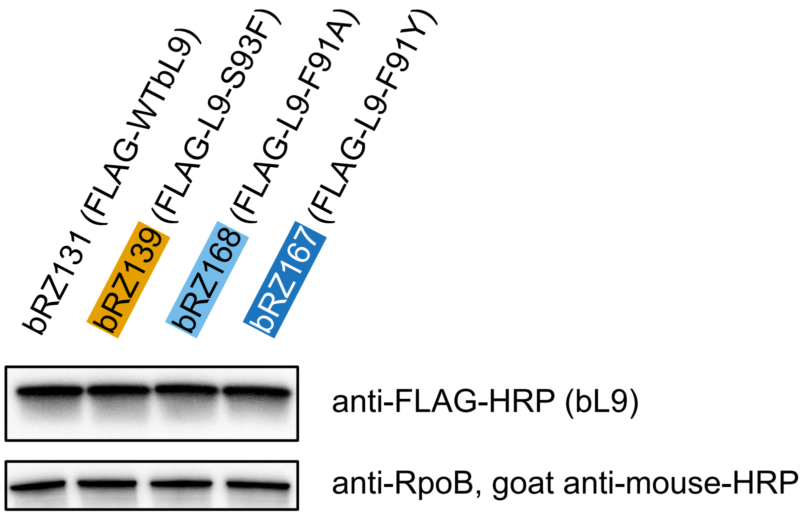


Figure S2. FLAG-bL9 expression levels are consistent

N-terminally FLAG-tagged bL9 variants were integrated at the genomic locus (marked with $Kan^R$). Cultures were grown in LB to midlog phase and whole cell lysates were prepared and normalized by BCA assay. An anti-FLAG (Sigma) Western blot was performed to compare expression levels of FLAG-bL9 between the strains. RpoB was used as a loading control.

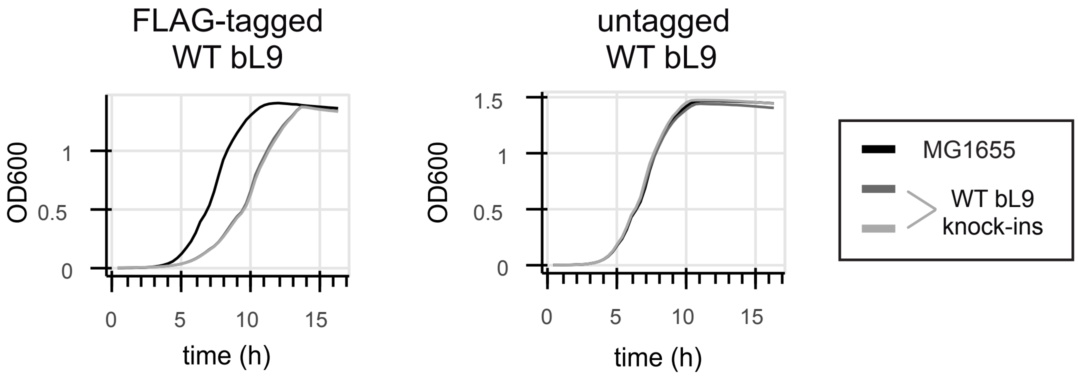


Figure S3. FLAG-tagged bL9 has a growth defect

FLAG-tagged or untagged wildtype bL9 was integrated at the genomic locus in a ΔL9 background strain (two separate lineages). Growth was compared to a reference wild type strain MG1655. Cells were diluted into a 96-well plate in LB + 50 µg/mL fusidic acid and grown at 37°C shaking in a plate reader; $OD_{600}$ readings were taken every 20 minutes. Each line is the average of replicate wells.

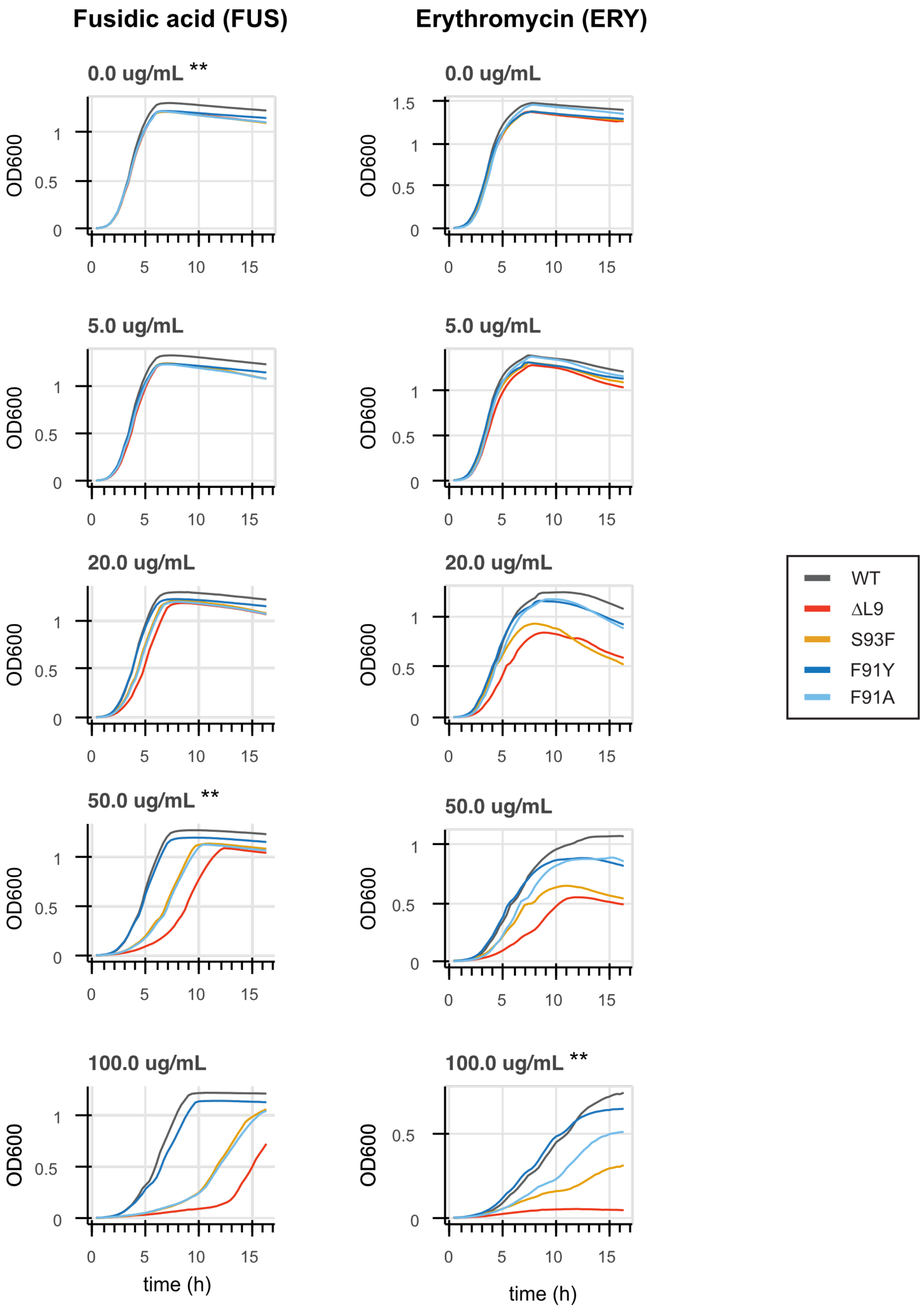


Figure S4. bL9 mutant growth curves in LB with various concentrations of fusidic acid and erythromycin. Each line is the average of replicate wells. This is a representative experiment, which was repeated three times. ** indicates panels that are included in Figure 2.

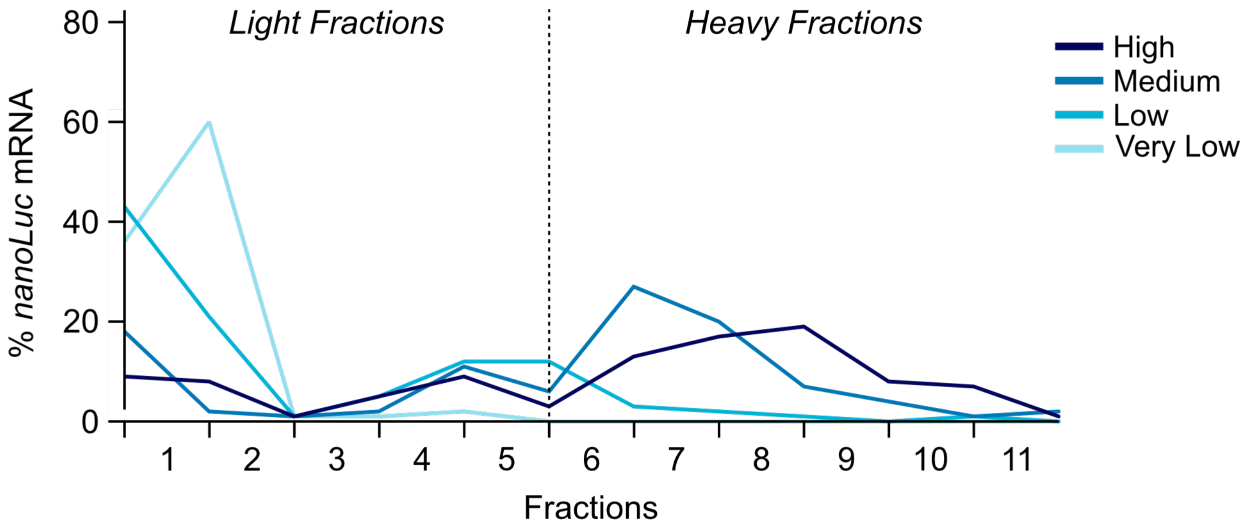


Figure S5. Distribution of *nanoLuc* mRNA within each fraction. The Shine Dalgarno reporter lysates were run on a 10-50% sucrose graident prior to RT-qPCR. This is a representative experiment, which was repeated three times.
